# Differential reaction norms to ocean acidification in two oyster species from contrasting habitats

**DOI:** 10.1101/2023.08.30.555611

**Authors:** Coline Caillon, Fabrice Pernet, Mathieu Lutier, Carole Di Poi

## Abstract

Ocean acidification (OA), a consequence of the increase in anthropogenic emissions of carbon dioxide, causes major changes in the chemistry of carbonates in the ocean with deleterious effects on calcifying organisms. The pH/pCO_2_ range to which species are exposed in nature is important to consider when interpreting the response of coastal organisms to OA. In this context, emerging approaches, which assess the reaction norms of organisms to a wide pH gradient, are improving our understanding of tolerance thresholds and acclimation potential to OA. In this study, we decipher the reaction norms of two oyster species living in contrasting habitats: the intertidal oyster *Crassostrea gigas* and the subtidal flat oyster *Ostrea edulis*, which are two economically and ecologically valuable species in temperate ecosystems. Six-month-old oysters of each species were exposed in common garden for 48 days to a pH gradient ranging from 7.7 to 6.4 (total scale). Both species are tolerant down to a pH of 6.6 with high plasticity in fitness-related traits such as survival and growth. However, oysters undergo remodelling of membrane fatty acids to cope with decreasing pH along with shell bleaching impairing shell integrity and consequently animal fitness. Finally, our work reveals species-specific physiological responses and highlights that intertidal *C. gigas* seems to have a better acclimation potential to rapid and extreme OA changes than *O. edulis*. Overall, our study provides important data about the phenotypic plasticity and its limits in two oyster species, which is essential for assessing the challenges posed to marine organisms by OA.

## INTRODUCTION

Since the Industrial Revolution, the increase in atmospheric CO_2_ emissions and its subsequent uptake by the ocean lead to an imbalance in seawater carbonate chemistry causing a decrease in pH, a process known as ocean acidification (OA) (Caldeira and Wickett, 2003). The most likely climate model predict that sea-surface pH will decrease by 0.032 pH units by the end of the century (SSP3-7.0; IPCC, 2022). Concomitantly, the excess of hydrogen ions (H^+^) is responsible for a reduction in the availability of carbonate ions 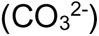, the essential components of the shells and skeletons of marine calcifiers (Orr et al., 2005).

In most experimental studies, marine organisms are exposed to constant acidification levels derived from the IPCC OA scenarios for the global ocean, while most of them inhabit variable coastal environments such as intertidal, estuarine, and upwelling areas (Cai et al., 2011; Feely et al., 2008; Provoost et al., 2010). In these habitats, current seawater pH/pCO_2_ levels vary more temporally and spatially than in the open ocean, and may already exceed average OA projections (Vargas et al., 2017). Therefore, this large environmental variability in combination with expected global future acidification could drive resident organisms even closer to the limits of their physiological tolerance (Hofmann et al., 2011). To meet this constraint, few studies have investigated the physiological tolerance and acclimation capacities of coastal organisms to OA by assessing reaction norms to a broad range of pH levels, covering the natural variability of seawater chemistry (Dorey et al., 2013; Lutier et al., 2022; Moulin et al., 2011; Thor et al., 2018; Ventura et al., 2016). In addition, reaction norms can be used to identify physiological tipping points, corresponding to tolerance thresholds beyond which a small change in an environmental factor has large effects on the organism. Such approaches have proven to be complementary methods to standard scenario studies (present-day vs. future), but they need to consider the entire natural pH/pCO_2_ range to which species are exposed to be relevant (Vargas et al., 2017).

Among coastal organisms, shelled molluscs are considered particularly vulnerable to OA as acidified conditions lead to a decrease in the calcite and aragonite saturation states, which adversely affects their calcification, shell integrity and survival (Gazeau et al., 2013; Waldbusser et al., 2015). Nevertheless, it is not worth noting that there are species- and population-specific responses to OA that can be explained by divergent selection and plasticity, which may or may not allow a set of organisms to cope with rapid and extreme acidification events (Duarte et al., 2015; Parker et al., 2010). Interestingly, molluscs living in environments with high natural pH/pCO_2_ variability would be less sensitive to OA (Thomsen et al., 2017; Vargas et al., 2017, 2022), and more specifically that intertidal organisms would have evolved greater tolerance to lower pH than their subtidal counterparts (Leung et al., 2017; Scanes et al., 2017).

Here, we aimed to determine and compare the reaction norms of two economically and ecologically valuable oyster species from different tidal habitats. The Pacific oyster *Crassostrea gigas* (Thunberg, 1793) predominantly inhabits the upper intertidal zone where they are exposed to severe metabolic hypercapnia during emersion (Burnett, 1988; Zwerschke et al., 2018). In contrast, the European flat oyster *Ostrea edulis* (Linnaeus, 1758) is mainly found in subtidal areas where they rarely or never emerge (Pogoda, 2019; Zwerschke et al., 2018). Therefore, we can expect the intertidal *C. gigas* to be more tolerant to pH decrease than its subtidal counterpart *O. edulis*.

Specifically, we determined the reaction norms of these two oyster species over a wide range of pH for survival and growth, two parameters that are related to animal fitness. We also investigated the effect of decreasing pH on shell coloration though quantitative image analysis. Indeed, acidification bleaches shells and the coloration could be a simple and reliable indicator of the acidity of water and its impact on the integrity of molluscs’ shell (Avignon et al., 2020). We also measured the energy reserves of oysters as a function of pH. Acclimation to acidification generally involves energy reallocation and strategies may differ depending on the species (Sokolova, 2013). Finally, we analysed the composition of membrane fatty acids, a key parameter governing exchanges between intra and extracellular compartments and overall metabolic rates (Hulbert, 2003). Overall, our study provides insight into phenotypic plasticity divergence in two oyster species from contrasting tidal habitats, and a better understanding of how oysters are likely to respond and acclimate in a rapidly acidifying ocean.

## MATERIAL AND METHODS

### Animals and maintenance

Due to different reproduction strategies between the two oyster species studied, the production and fertilisation protocols used were different. Pacific oysters were produced at the Ifremer experimental station (Argenton, Brittany, France) on 27 August 2019, according to the procedure described in Petton et al. (2015). Briefly, the broodstock consisted of 118 females and 22 males collected in the natural environment between 2011 and 2017 in Île d’Aix (Charente-Maritime, France), whose gametes were obtained by stripping and mixed in the same jar to fertilise the oocytes. At one-month-old, the juvenile oysters were transferred to the Ifremer station in Bouin (Vendée, France) for growing. On 9 January 2020, these oysters were brought back to the Argenton facilities and kept in a 500 l flow-through tank for 19 days before the experiment.

In the flat oyster, the fertilisation is internal and the females incubate the larvae for nearly 10 days in the paleal cavity. Compared with the Pacific oyster, hatchery culture of the flat oyster is delicate and mortality occurs (Maneiro et al., 2020). We used juvenile oysters obtained by biparental mating (paired crossing) at the hatchery of the Comité Régional de la Conchyliculture (Lampaul-Plouarzel, Brittany, France) on 19 August 2019. The two parents used for breeding were wild oysters collected in the Bay of Brest in May 2019. On 17 January 2020, the juvenile oysters were transferred to the Argenton facilities and kept for 11 days in the same flow-through tank as the Pacific oysters.

During this time, the seawater temperature was gradually increased from 14°C to 19°C (+0.5°C.day^-1^), which is an optimal temperature for both species (Bayne, 2017). Oysters were continuously fed a mixed diet of two phytoplankton species (1:1 in dry weight), the diatom *Chaetoceros muelleri* (strain CCAP 1010/3) and the haptophyte *Tisochrysis lutea* (strain CCAP 927/14). Phytoplankton concentration was maintained at 1500 μm^3^.µl^-1^ at the tank outlet to ensure *ad libitum* feeding (Petton et al., 2015), and monitored twice daily using an electronic particle counter (Multisizer 3, Beckman Coulter, Indianapolis, USA; 100-µm aperture tube). On 28 January 2020, at the onset of the experiment, the mean individual shell length and total mass were 18.5 ± 3.0 mm and 0.67 ± 0.18 g for *C. gigas*, and 8.9 ± 2.1 mm and 0.10 ± 0.08 g for *O. edulis*. Both species were divided into 16 batches containing 105 ± 3 *C. gigas* and 105 ± 4 *O. edulis* for a total biomass of 91.9 ± 2.4 g.

### Experimental design

Each batch of oysters was exposed in common garden to one constant nominal total pH (pH_T_) condition ranging from pH 7.9 to 6.4 with a step of 0.1 between two levels, at 19°C for 48 days and fed *ad libitum* as previously described. The experimental system consisted of 17 experimental units that were randomly assigned to one pH condition (n = 16) or to a control blank without animals (n = 1) to check the pH of the ambient seawater. Each experimental unit consisted of a header tank in which seawater was acidified by injecting pure CO_2_ (except for the ambient pH) and then delivered by a pump (ProFlow t500; JBL, Neuhofen, Germany) to a holding tank containing the oysters. These tanks were 45 l and their entire volume was renewed every 90 min.

The pH level was controlled in the header tank using a pH-controller connected to a pH electrode (ProFlora; JBL, Neuhofen, Germany). The pH electrode was calibrated with NBS (National Bureau of Standards) buffers (pH 4.00 and 7.00) at the beginning of the experiment. In the holding tank, the seawater was continuously oxygenated and mixed using air bubbling and a circulation pump (ProFlow t300; JBL, Neuhofen, Germany). The tanks and oysters were cleaned twice a week. The photoperiod was fixed at 14 light: 10 dark and no tidal regime was applied. Seawater was sampled twice daily at the inlet and outlet of each holding tank to determine the phytoplankton concentration (in cell volume) and total consumption of both species simultaneously using the electronic particle counter. When needed in each pH condition, the phytoplankton concentration at the inlet was gradually increased over time to compensate for the increasing grazing rate of the growing oysters (Fig. S1). Considering the seawater renewal time of the tanks, pH had no effect on the quality of microalgae (Fig. S2).

### Seawater carbonate chemistry

Seawater pH_T_, temperature, dissolved oxygen saturation and salinity were monitored twice daily in the holding tanks using a portable multi-parameter device (MultiLine® Multi 3630 IDS - WTW: pH electrode SenTix® 940, oxygen sensor FDO® 925, conductivity electrode TetraCon® 925; Xylem Analytics, Weilheim in Oberbayern, Germany). These measurements were used to adjust the settings of the pH-controllers. The accuracy of the pH electrode was checked once a week with Certipur® NBS buffers (pH 4.00, 7.00 and 9.00; Merck, Darmstadt, Germany) and calibrated twice a week on the total scale (pH_T_) with a certified Tris/HCl buffer at a salinity of 35.0 (provided by A. G. Dickson, Scripps Institution of Oceanography, San Diego, USA). Seawater samples (150 ml) were collected three times during the experiment for total alkalinity (A_T_) analyses. The seawater was filtered through 0.7 µm GF/F glass microfiber filters (Whatman®, Florham Park, USA) and immediately poisoned with 0.05% saturated mercuric chloride solution before storage. A_T_ was measured at the Laboratoire Environnement Ressources (Sète, Hérault, France) using potentiometric titrations with an automatic titrator (TitroLine® 7000; SI Analytics, Mainz, Germany) according to Dickson et al. (2007). A_T_ measurements were conducted in triplicate at 20°C on 25 ml subsamples with a measurement uncertainty of 0.44%. Parameters of the carbonate system were determined from pH_T_, A_T_, temperature and salinity using the R package *seacarb* (v3.3.1; Gattuso et al., 2022).

### Oyster survival

For each species, oyster survival was assessed once daily. The dead individuals were immediately removed.

### Biometric measurements

Total body weight (shell + tissue), and both shell length and weight, were measured individually on a subsample of 30 individuals of each species at the start of the experiment (day 0), and on 20 oysters per species and pH condition after 41 days of exposure. The dissected flesh was pooled and weighted to obtain fresh flesh weight. Growth rate (G) was calculated as:

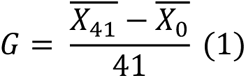

where G is the growth rate expressed as increase in shell length or total body weight per day (mm.d^−1^ and mg.d^−1^, respectively), 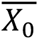 and 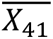 are the mean values measured at the onset of the experiment (day 0) and at 41 days. The measurements relate to different individuals between day 0 and day 41.

### Shell coloration

Analysis of shell coloration was performed on day 20, as shell condition deteriorated under the lowest pH conditions afterwards. The digestive gland was apparent through the shell later in the exposure and could distort shell coloration measurements. The left valves of 15 individuals per species and pH condition were randomly selected. The shells were photographed using an ultra-high precision digital microscope (Keyence® VHX-5000; Keyence Corporation, Osaka, Japan). The images were converted into grayscale images and analysed using ImageJ software. The mean coloration of the shell was calculated using the mean value of the grey levels of all pixels constituting the selected area (Fig. 1). Grayscale level was measured on a scale of 0-255 where 0 represents black and 255 represents white.

**Fig. 1.**
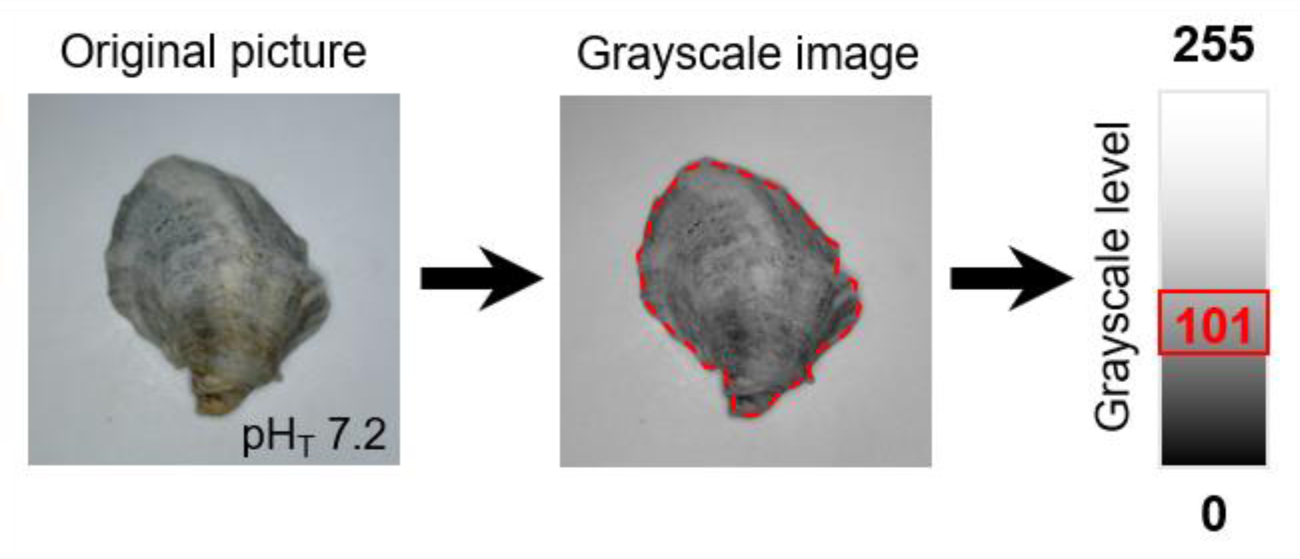
Schematic representation of the procedure for analysing the coloration of oyster shells. The shell coloration was determined using ImageJ software that calculated the mean grayscale level of the entire left shell valve ranging from 0 (total black) to 255 (total white).

### Biochemical analyses

Soft-tissue samples from 10 individuals per species and pH condition were collected at day 41, flash-frozen under liquid nitrogen, pooled, powderised with a ball mill and stored at −80°C until analyses. Lipids and carbohydrates were not analysed for flat oysters exposed to pH below 6.8 and 6.9, respectively, due to the low amount of flesh available.

#### Energy reserves

Oyster powder was diluted with chloroform/methanol (2:1, v/v) and the neutral lipids (i.e., reserve lipids) were separated into lipid classes and quantified using a high-performance thin-layer chromatography and a scanning densitometer (Automatic TLC Sampler 4 and TLC Scanner 3, respectively; CAMAG, Muttenz, Switzerland). Since triacylglycerols (TAG) are mainly reserve lipids and sterols (ST) are structural lipids of cell membranes, the TAG:ST ratio was used as an index for the contribution of energy reserve to structure (Fraser, 1989).

The carbohydrate content was determined according to the colorimetric method described in Dubois et al. (1956). Milli-Q water was added to the powder samples and mixed with phenol (0.5 ml, 5% m/v) and sulphuric acid (2.5 ml, 98%), and then incubated for 40 min. Absorbance was measured at 490 nm using a Cary 60 UV-Vis spectrophotometer (Agilent Technologies, Santa Clara, USA). Total carbohydrate content was calculated using a standard calibration curve with pure glucose and expressed in mg.g^-1^ fresh weight.

Proteins were extracted from powder on ice using lysis buffer [150 mM NaCl (Merck), 10 mM Tris/HCl (Sigma-Aldrich), 1 mM EDTA (Quantum), 1 mM EGTA (Sigma-Aldrich), 1% Triton X-100 (Bio-Rad), and 0.5% Igepal (Sigma-Aldrich); pH 7.4 at 4°C] with phosphatase and protease inhibitors [1% of phosphatase inhibitor cocktail II (Sigma-Aldrich), 2% NaPPi 250 mM (Sigma-Aldrich), and a tablet of complete EDTA free protease inhibitor cocktail (Roche) in 25 mL of lysis buffer]. Samples were ground and homogenised using a Polytron® PT 2500 E (Kinematica, Malters, Switzerland). The resulting lysates were used for quantification of the total protein content with the DC protein assay (Bio-Rad, Hercules, USA) according to Lowry et al. (1951). Absorbance was read at 750 nm and protein concentrations were determined by comparison with a standard calibration curve supplied with the kit. The results were expressed as mg.g^-1^ fresh weight.

#### Fatty acid composition of polar lipids

Polar lipids (i.e., membrane lipids) were purified from the chloroform/methanol mixture used for lipid class analysis on a silica-gel micro-column, and transesterified with a mixture of 3.4% sulphuric acid H_2_SO_4_ in methanol at 100°C (Couturier et al., 2020). This transesterification produces fatty acid methyl esters (FAME) from the fatty acids, and dimethyl acetals (DMA) from the alkenyl chains at the sn-1 position of plasmalogens. The resulting FAME and DMA were analysed using a flame ionization detector gas chromatograph system (HP - Agilent 6890; Agilent Technologies, Santa Clara, USA) equipped with a DB-Wax capillary column (30 m length x 0.25 mm i.d. x 0.25 µm film thickness). The DMA quantification allowed the indirect quantification of plasmalogens. Fatty acids (FA) were then identified by comparing of retention times with standards. Each FA was expressed as the relative percentage of peak area by the sum of all polar FA peaks.

### Statistical analyses

All statistical analyses were performed using RStudio software (v4.2.2) and the significance threshold was set at 0.05. All variables were plotted using the mean pH_T_ calculated from daily measurements throughout the experiment in each pH condition. The general statistical procedure was previously described in Lutier et al. (2022).

The relationships between dependent variables (survival rate, biometrics, shell coloration, biochemistry) and the mean pH_T_ were computed using regression models. Segmented and linear regression models were tested and compared for each variable. The model with the lowest Akaike and Bayesian information criteria (AIC, BIC) and the highest R^2^ was selected. For segmented regression models, both the tipping point and its 95% confidence interval were estimated. The tipping point corresponds to the pH value at which the dependent variable tips, and was defined by applying the bootstrap restarting algorithm (Wood, 2001). For the growth rates, the critical point – i.e., the pH value at which the dependent variable was zero – and its 95% confidence interval were also determined. The normality of residuals and homogeneity of variances were graphically checked. The significance of each slope was tested using the Student t-test and compared to a horizontal slope of zero. The segmented regression models were computed using *segmented* package (v1.6-2). Dependent variables were averaged over the number of individuals collected for each species and pH condition.

In addition, slopes of each regression model obtained for *C. gigas* and *O. edulis* were compared using the Welch two-sample t-test to detect significant differences in response to acidification between the two species. The comparison of slopes was carried out above and below the tipping point when one of the species’ reaction norms has been modelled by a segmented model. The test was not performed if the slopes were not different from zero. The statistical outputs are summarised in Table 2.

Specifically, survival of both species was evaluated by the Kaplan-Meier (KM) method (Kaplan and Meier, 1958) for each pH condition. Survival time was measured in days from the onset of the experiment. KM survival curves were generated using the *survival* package (v3.5-5). Multiple pairwise comparisons were then made using the Log-Rank survival test to identify significant differences in survival between the different pH conditions. Final survival rates were extracted and tested against pH_T_.

Fatty acid data were summarised using principal component analysis (PCA) for each species separately. Only fatty acids contributing to >1% were considered. The pH condition 7.5 for *C. gigas* was removed from the fatty acid data analysis due to extraction problems. The fatty acids were then divided into two groups according to their positive or negative correlation with the first principal component (PC1). The contribution of fatty acids (%) was then summed for each group and plotted as a function of pH_T_.

## RESULTS

### pH regulation and carbonate chemistry

During the experiment, pH levels in the oyster tanks were stable and reached the targeted values, except for the highest pH conditions that was 7.7 instead of 7.8 and 7.9 (Table 1). This was due to oyster respiration, which lowered the pH of the incoming seawater (already at a pH_T_ of 7.9), despite a relatively high turnover rate of seawater relative to oyster biomass. Total alkalinity (A_T_) varied from 2321 to 2458 μmol.kg^−1^ and increased significantly with decreasing pH_T_ (t-test, p < 0.001, R^2^ = 0.95) up until reaching control values (seawater with no oyster). Seawater was undersaturated (Ω < 1) with respect to aragonite and calcite from pH levels 7.5 and 7.3, respectively.

**Table 1.** Parameters of seawater carbonate chemistry in each experimental tank (16 pH_T_ levels and a blank without oysters).

| Nominal |  | Measured |  |  |  | Calculated |  |  |  |
| --- | --- | --- | --- | --- | --- | --- | --- | --- | --- |
| pH | $\text{pH}_T$ | $A_T$<br>($\mu\text{mol.kg}^{-1}$) | T<br>($^{\circ}\text{C}$) | S<br>(ppt) | DO<br>(%) | $\text{pCO}_2$<br>($\mu\text{atm}$) | DIC<br>($\mu\text{mol.kg}^{-1}$) | $\Omega_C$ | $\Omega_A$ |
| Blank | 7.9±0.0 | 2444±6 | 19.1±0.1 | 34.6±0.2 | 100.4±0.5 | 693±22 | 2275±8 | 3.2±0.1 | 2.1±0.0 |
| 7.9 | 7.7±0.0 | 2349±29 | 19.1±0.1 | 34.6±0.2 | 98.7±0.7 | 947±18 | 2239±23 | 2.3±0.1 | 1.5±0.1 |
| 7.8 | 7.7±0.0 | 2350±54 | 19.2±0.1 | 34.6±0.2 | 98.6±0.6 | 964±54 | 2242±44 | 2.3±0.2 | 1.5±0.1 |
| 7.7 | 7.7±0.0 | 2321±37 | 19.2±0.1 | 34.6±0.2 | 98.5±0.7 | 1085±51 | 2233±39 | 2.0±0.0 | 1.3±0.0 |
| 7.6 | 7.6±0.1 | 2339±57 | 19.1±0.1 | 34.6±0.2 | 97.7±0.9 | 1382±312 | 2281±79 | 1.7±0.3 | 1.1±0.2 |
| 7.5 | 7.5±0.0 | 2345±27 | 19.1±0.1 | 34.6±0.2 | 98.6±0.6 | 1814±59 | 2325±29 | 1.3±0.0 | 0.9±0.0 |
| 7.4 | 7.4±0.0 | 2368±55 | 19.1±0.1 | 34.6±0.2 | 98.4±0.7 | 2292±126 | 2378±57 | 1.1±0.0 | 0.7±0.0 |
| 7.3 | 7.3±0.0 | 2361±29 | 19.1±0.1 | 34.6±0.2 | 98.1±0.9 | 2901±575 | 2403±55 | 0.9±0.1 | 0.6±0.1 |
| 7.2 | 7.2±0.0 | 2369±28 | 19.1±0.1 | 34.6±0.2 | 98.2±0.7 | 3405±153 | 2436±22 | 0.8±0.0 | 0.5±0.0 |
| 7.1 | 7.1±0.0 | 2391±33 | 19.2±0.1 | 34.6±0.2 | 98.8±0.6 | 4578±125 | 2508±37 | 0.6±0.0 | 0.4±0.0 |
| 7.0 | 7.0±0.0 | 2412±60 | 19.2±0.1 | 34.6±0.2 | 98.7±0.6 | 5759±152 | 2574±59 | 0.5±0.0 | 0.3±0.0 |
| 6.9 | 6.9±0.1 | 2404±22 | 19.1±0.1 | 34.6±0.2 | 98.8±0.5 | 6808±476 | 2606±4 | 0.4±0.0 | 0.3±0.0 |
| 6.8 | 6.8±0.1 | 2426±20 | 19.2±0.1 | 34.6±0.2 | 99.2±0.4 | 9655±1173 | 2728±59 | 0.3±0.0 | 0.2±0.0 |
| 6.7 | 6.7±0.1 | 2433±21 | 19.1±0.1 | 34.6±0.2 | 99.0±0.5 | 10482±631 | 2765±23 | 0.3±0.0 | 0.2±0.0 |
| 6.6 | 6.6±0.0 | 2445±15 | 19.2±0.1 | 34.6±0.2 | 99.1±0.4 | 15930±2494 | 2962±93 | 0.2±0.0 | 0.1±0.0 |
| 6.5 | 6.5±0.1 | 2458±23 | 19.1±0.1 | 34.6±0.2 | 98.9±0.6 | 20108±1915 | 3117±70 | 0.1±0.0 | 0.1±0.0 |
| 6.4 | 6.4±0.0 | 2453±24 | 19.1±0.1 | 34.6±0.2 | 98.7±0.6 | 22186±1493 | 3182±72 | 0.1±0.0 | 0.1±0.0 |

Seawater pH on the total scale ( $\text{pH}_T$ ), temperature (T), dissolved oxygen saturation (DO) and salinity (S) were measured twice daily and averaged over the entire experimental period, while total alkalinity ( $A_T$ ) was measured three times over the experiment. Partial pressure of $\text{CO}_2$ ( $\text{pCO}_2$ ), dissolved inorganic carbon concentration (DIC), saturation state of calcite ( $\Omega_C$ ) and aragonite ( $\Omega_A$ ) were calculated from $\text{pH}_T$ , $A_T$ , S and T using the R package *seacarb* (Gattuso et al., 2022). Values correspond to mean ± s.d.

### Low pH induces oyster mortality particularly for *O. edulis*

No mortality was observed in *C. gigas* throughout the exposure period, except at the lowest pH_T_ 6.4 where mortalities started at day 38 and survival reached 78% at the end of the experiment (Fig. 2A). No pH-based model fitted with the final survival of *C. gigas* (Fig. 2C). In contrast, in *O. edulis*, mortalities occurred in all pH conditions throughout the exposure period, and survival reached 38% at the lowest pH value (Fig. 2B). The final survival of *O. edulis* fitted with pH according to a segmented regression model with a tipping point at pH_T_ 6.6 below which survival declined drastically with decreasing pH (Fig 2C). Above this tipping point, survival ranged from 93% to 75% and did not correlate with pH (t-test, p = 0.98).

**Fig. 2.**
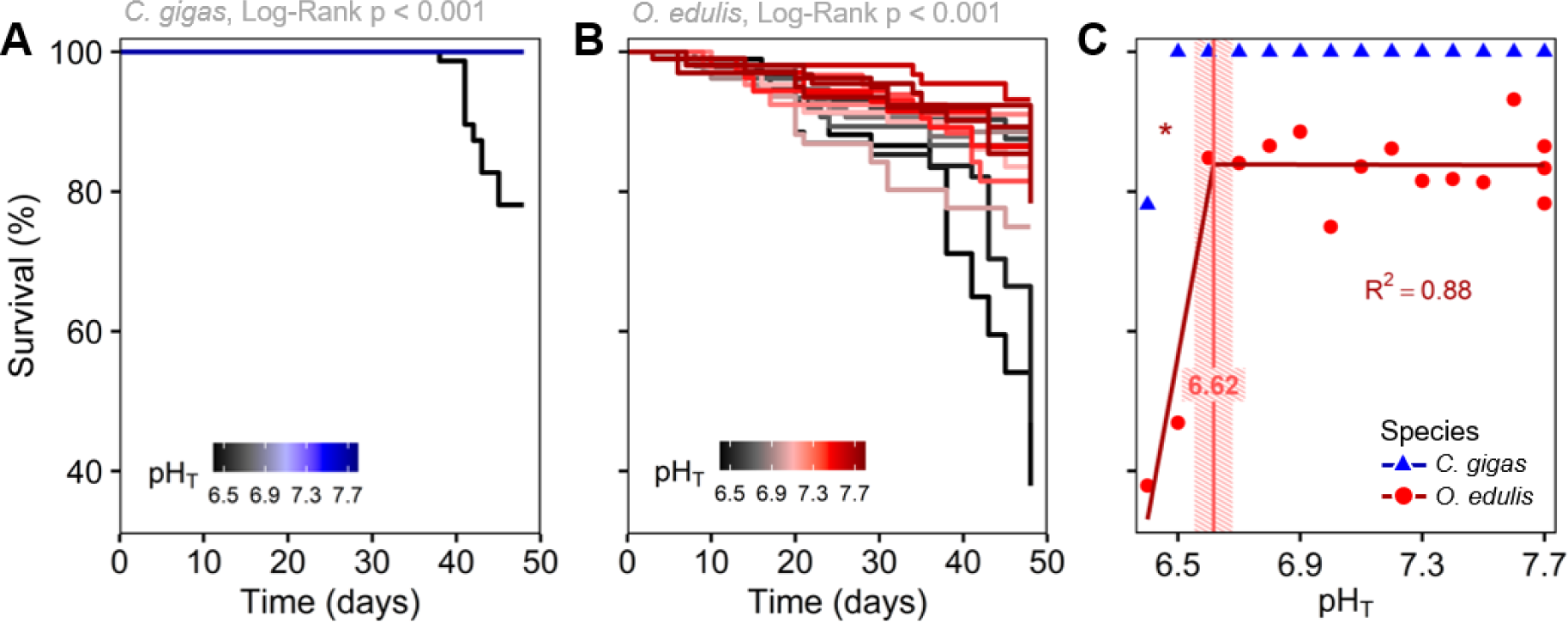
Survival of oysters for each species as a function of exposure time and pH (on total scale). Kaplan-Meier survival curves of *Crassostrea gigas* (A) and *Ostrea edulis* (B) oysters during 48 days of exposure under 16 pH_T_ conditions, and final survival of oysters tested against pH_T_ (C). Tipping point and the 95% confidence interval for survival in *O. edulis* are shown in striped red, not applicable for *C. gigas*. The significance level of slope is represented using symbols (p <0.05*, <0.1·).

### Growth declines with decreasing pH more markedly for *C. gigas*, and shell dissolution occurred at pH 6.6 for both species

At day 41, all biometric parameters declined with decreasing pH for both oyster species (Fig. 3). In *C. gigas*, shell length and growth rates (based on shell length and flesh weight) decreased significantly as the pH decreased, and reached a tipping point at pH_T_ ∼7.2 below which they declined sharply (Fig. 3A, B and F). In contrast, shell weight, total body weight and growth rate calculated from body weight decreased linearly over the entire pH range (Fig. 3C, D and E). In *O. edulis*, all parameters decreased linearly with pH and showed no tipping point (Fig. 3A-F). Growth rates (calculated from shell length and shell weight) were negative when pH was lower than 6.6 (i.e., critical points) for both species (Fig. 3B and E). For most parameters, slope comparison tests on either side of the tipping point indicated that the reaction norms to pH differed significantly between the two species, and that *C. gigas* generally showed steeper slopes (Table 2).

**Table 2.** Comparison of the reaction norms for whole-body and shell-related parameters in *Crassostrea gigas* and *Ostrea edulis* in response to ocean acidification.

| Variable | Species | Model results |  | Slope value |  | Slope comparison |  |
| --- | --- | --- | --- | --- | --- | --- | --- |
|  |  | Regression | TP | <TP | >TP | <TP | >TP |
| Shell length | *C. gigas* | segmented | 7.20±0.28 | 16.97±1.93 | 6.64±2.41 | <0.001 | 0.32 |
|  | *O. edulis* | linear | n.a. | 7.55±0.61 |  |  |  |
| Total body weight | *C. gigas* | linear | n.a. | 2.29±0.14 |  | <0.001 |  |
|  | *O. edulis* | linear | n.a. | 0.50±0.04 |  |  |  |
| Shell weight | *C. gigas* | linear | n.a. | 1.55±0.09 |  | <0.001 |  |
|  | *O. edulis* | linear | n.a. | 0.40±0.03 |  |  |  |
| Flesh weight | *C. gigas* | segmented | 7.16±0.26 | 0.55±0.06 | 0.21±0.07 | <0.001 | 0.001 |
|  | *O. edulis* | linear | n.a. | 0.07±0.01 |  |  |  |
| Growth rate (length) | *C. gigas* | segmented | 7.20±0.28 | 0.41±0.05 | 0.16±0.06 | <0.001 | 0.32 |
|  | *O. edulis* | linear | n.a. | 0.18±0.01 |  |  |  |
| Growth rate (weight) | *C. gigas* | linear | n.a. | 55.85±3.39 |  | <0.001 |  |
|  | *O. edulis* | linear | n.a. | 12.16±0.98 |  |  |  |
| Shell coloration | *C. gigas* | linear | n.a. | -32.57±4.13 |  | / | <0.001 |
|  | *O. edulis* | segmented | 6.93±0.25 | n.s. | -44.77±7.44 |  |  |

Comparison of the slopes was performed above and below the tipping point when one of the species displayed a segmented model. The test was not performed on slopes not significantly different from a zero horizontal slope. Values are TP or slope coefficient and their 95% confidence interval. Welch two-sample t-test was performed to compare species slopes and significant p-values ( $p < 0.05$ ) are indicated in bold. Abbreviations: TP, tipping point; <TP, below tipping point; >TP, above tipping point.

**Fig. 3.**
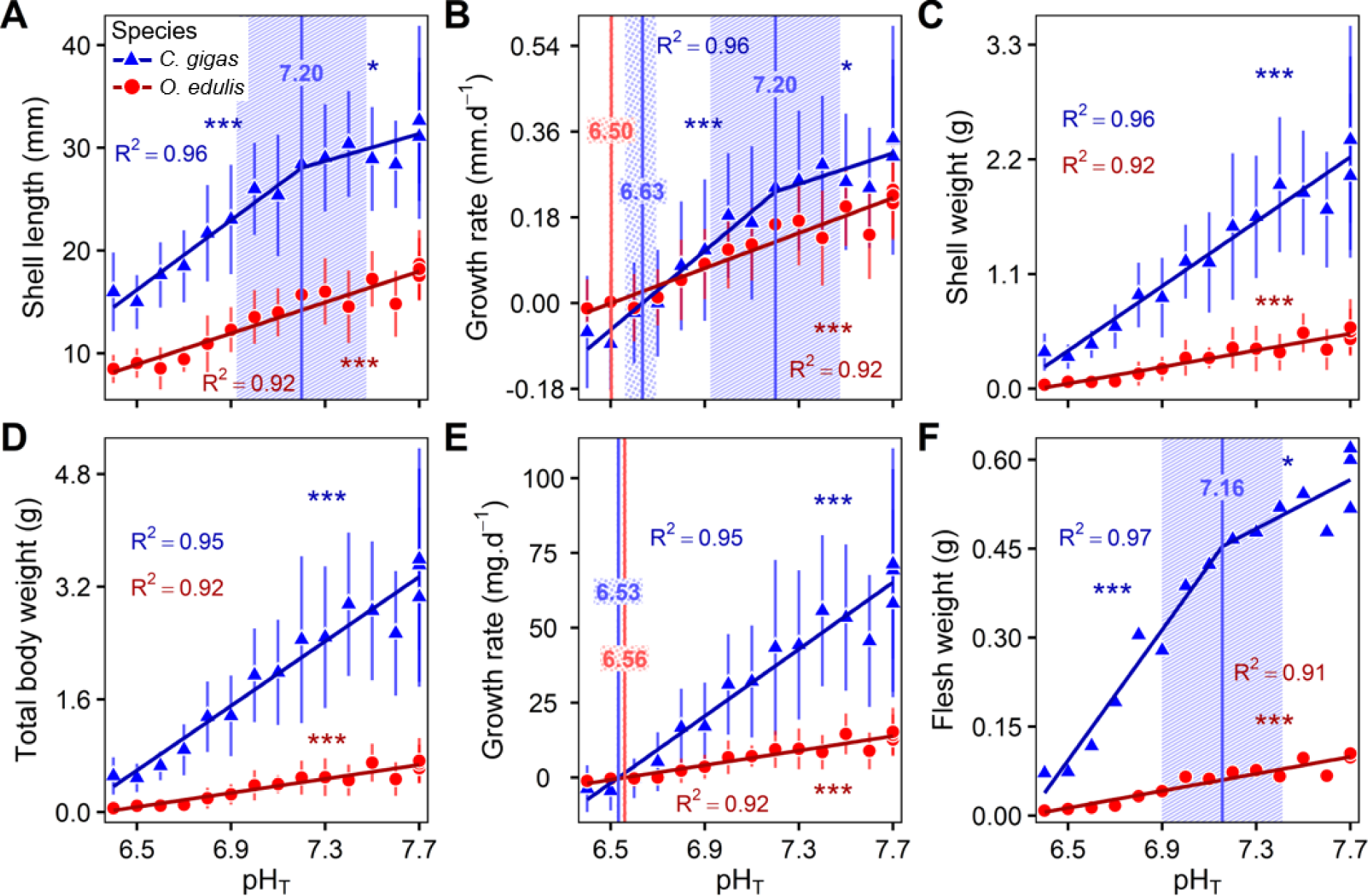
Final biometric parameters in *C. gigas* and *O. edulis* oysters as a function of pH (on total scale). Shell length (A), growth rate in shell length (B), shell weight (C), total body weight (D), growth rate in total body weight (E) and fresh flesh weight (F) in oysters after 41 days of exposure under 16 pH_T_ conditions. Data correspond to mean ± s.d. when available (n = 20 per pH condition and species). Tipping points and the 95% confidence intervals are shown in striped blue for *C. gigas* and striped red for *O. edulis*. Critical points and the 95% confidence intervals are shown in dotted blue for *C. gigas* and dotted red for *O. edulis*. The significance level of slopes is represented using symbols (p <0.001***, <0.01**, <0.05*, <0.1·).

### Shell bleaching increases with decreasing pH for both species

After 20 days of exposure, the mean grayscale value increased with decreasing pH for both species. This indicates that decreasing pH increased shell bleaching (Fig. S3). Shell coloration changed linearly over the pH range in *C. gigas*, while it tipped at pH 6.9 in *O. edulis* and below which it plateaued (t-test, p = 0.58; Fig. 4). The grayscale values of the shell were similar between the two species at pH_T_ 7.7 (F = 13.1, p = 0.07). Slope comparison test (above the tipping point) indicated that *O. edulis* showed the steepest slope for shell bleaching (Table 2).

**Fig. 4.**
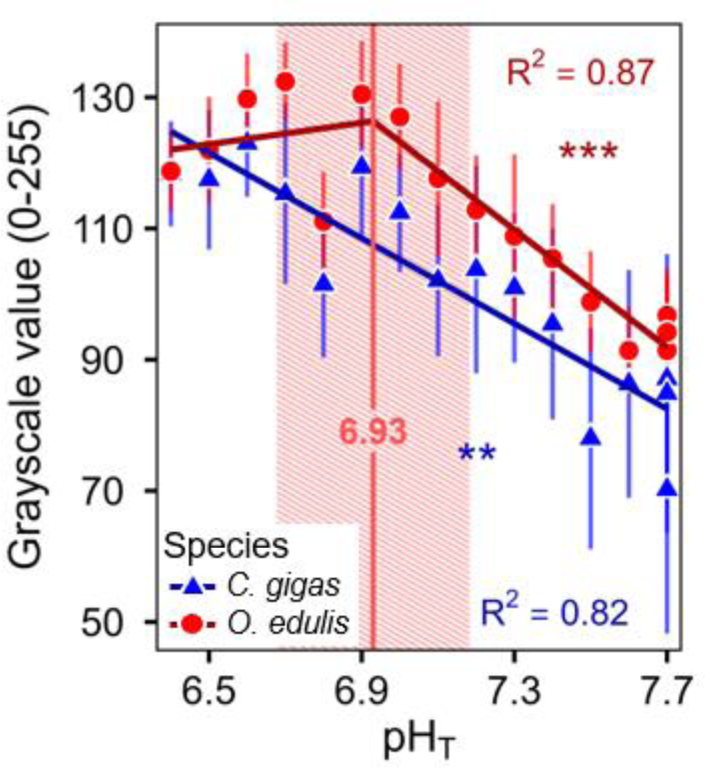
Shell coloration in *C. gigas* and *O. edulis* oysters as a function of pH (on total scale). Grayscale value corresponds to the shell coloration ranging from 0 (total black) to 255 (total white), after 20 days of exposure under 16 pH_T_ conditions. Data correspond to mean ± s.d. (n = 15 per pH condition and species). Tipping points and the 95% confidence intervals are shown in striped red for *O. edulis*. The significance level of slopes is represented using symbols (p <0.001***, <0.01**, <0.05*, <0.1·).

### Remodeling of membrane lipids occurs in both oyster species in response to decreasing pH, but involves different fatty acids and different tipping points

Principal component analysis of membrane fatty acids showed that the first axis (PC1) alone explained 61% and 47% of the total variance in relation to pH for *C. gigas* and *O. edulis* respectively (Figure 5A-B). Positively correlated fatty acids mainly consisted of palmitic acid (PA, 16:0), docosahexaenoic acid (DHA, 22:6n-3) and docosapentaenoic acid (DPA, 22:5n-6) in *C. gigas* (Table S1), while they were DHA, DPA and monounsaturated FA (18:1n-7) in *O. edulis* (Table S2). This positively correlated group of FAs exhibited tipping points at pH_T_ ∼ 7.0 for *C. gigas* and pH_T_ ∼ 7.5 for *O. edulis*, below which their contribution to membranes decreased significantly (Fig. 5C). Concomitantly, the main negatively correlated FAs consisted of non-methylene-interrupted FA (22:2NMI_i,j_), monounsaturated FA (20:1n-7) and eicosapentaenoic acid (EPA, 20:5n-3) in *C. gigas* (Table S1), while they were dimethyl acetal (18:0DMA) and stearic acid (SA, 18:0) in *O. edulis* (Table S2). This negatively correlated group of FAs exhibited tipping points at pH_T_ ∼ 7.0 for *C. gigas* and pH_T_ ∼ 7.5 for *O. edulis*, below which their contribution to membranes increased significantly (Fig. 5D).

**Fig. 5.**
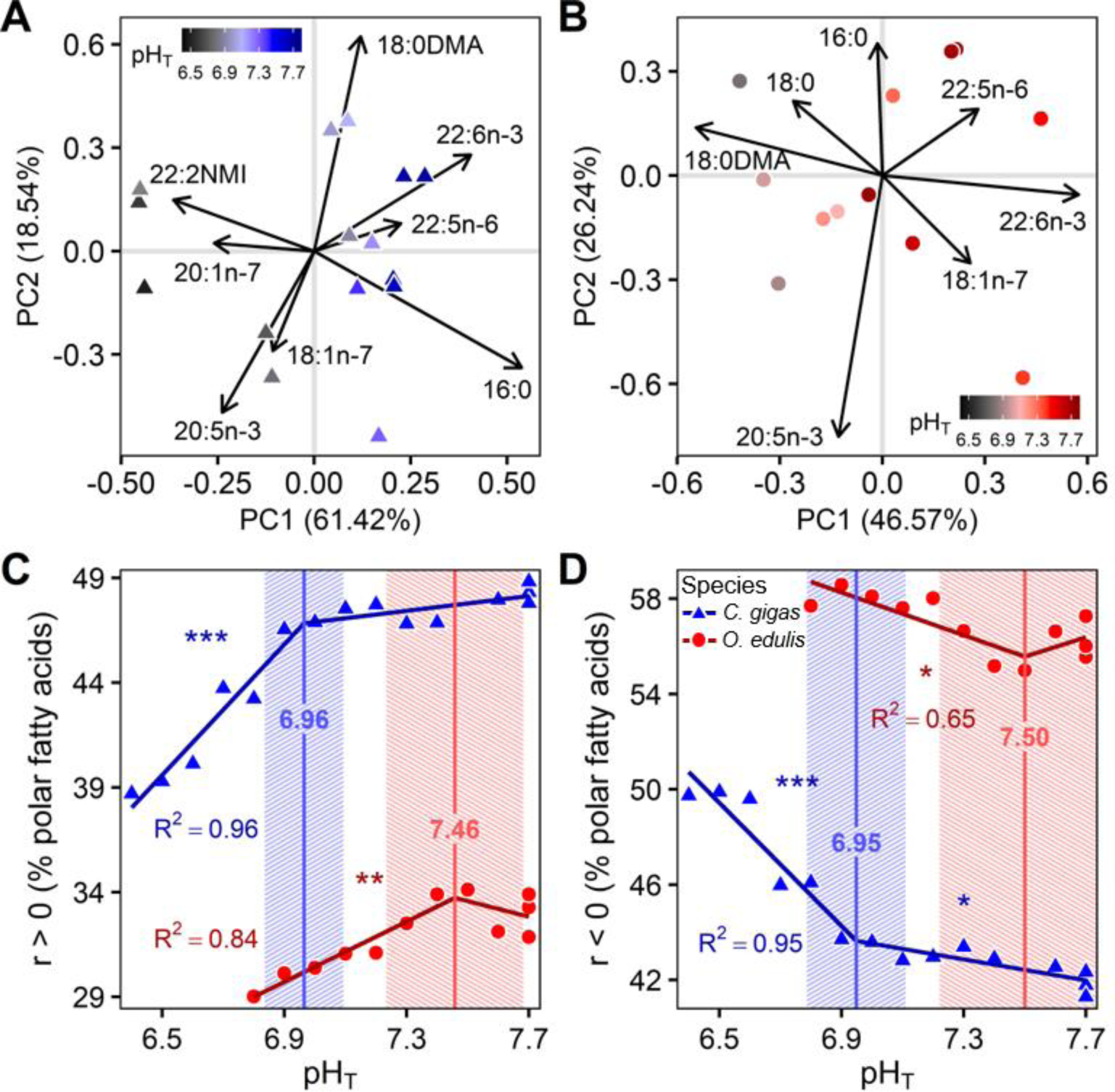
Membrane fatty acids (FA) composition in *C. gigas* and *O. edulis* oysters as a function of pH (on total scale). Principal component analysis of polar FA of *C. gigas* (A) and *O. edulis* (B) after 41 days of exposure under 16 pH_T_ conditions (n = 10 oysters pooled per pH condition and species). Arrows represent FA contributing to more than 5% of the first and second principal components (PC1 and PC2). The contributions of FA positively (C) or negatively (D) correlated to PC1 were summed together and tested against pHT for each oyster species. Tipping points and the 95% confidence intervals are shown in striped blue for *C. gigas* and striped red for *O. edulis*. The significance level of slopes is represented using symbols (p <0.001***, <0.01**, <0.05*, <0.1·).

### Energy reserves were unaffected by decreasing pH for both species

No relationship was found between TAG:ST ratio, total carbohydrate and total protein contents and pH_T_ for both oyster species (Fig. S4), although a segmented regression model was marginally significant for total proteins in *O. edulis* (t-test below tipping point, p = 0.08; Fig. S4C).

## DISCUSSION

Here, we determined and compared the reaction norms to OA in two oyster species living in different habitats: the intertidal Pacific oyster *C. gigas* and the subtidal European flat oyster *O. edulis* exposed in common garden to a seawater pH_T_ from 7.7 to 6.4 for 48 days. We established the reaction norms of juvenile oysters for survival rate, biometrics, shell coloration, membrane fatty acids and energy reserves vis-à-vis pH_T_. We found that both species showed high plasticity since they were able to survive and grow at extremely low pH_T_ (> 6.6). However, we found that lowering the pH decreased growth performance, increased shell bleaching and induced a major remodeling in membrane lipids. Interestingly, we found several interspecific differences, which suggest that the intertidal Pacific oyster has a higher acclimation potential than the subtidal flat oyster to cope with rapid and extreme acidification events.

### OA reduces growth-related parameters and causes shell bleaching of oysters

Growth-related parameters decreased as soon as pH declined in both oyster species, probably reflecting the lowering of the calcium carbonate (CaCO_3_) saturation state, and/or the increase in the metabolic cost of maintaining homeostasis (Beniash et al., 2010; Cyronak et al., 2016). The tipping points observed in *C. gigas* on the biometric parameters in this study are similar to those observed by Lutier et al. (2022) who used the same experimental design. Nevertheless, we obtain significant slopes above tipping points that Lutier et al. did not observe. This difference in pH responses between the two studies may be associated with differences in the experimental conditions or in the oyster populations used. Indeed, seawater temperature was 19°C in our study compared to 22°C in Lutier et al. (2022). Higher temperature can mitigate the impacts of a pH reduction (Di Poi et al., 2022; Ko et al., 2014). Besides, we used oysters from the same origin as Lutier et al. (2022) but in different years, so their life history and their susceptibility to acidification may have been different (Duarte et al., 2015; Parker et al., 2011; Thor et al., 2018). There are no comparable studies for *O. edulis* in the literature. Overall, the negative impacts on growth-related parameters as soon as pH falls could affect animals’ fitness, raising concerns about the future of oyster populations with even small changes in ocean acidity.

In addition, we found that shell bleaching increased with decreasing pH for both species. *O. edulis* exhibited a tipping point at pH_T_ 6.9 below which the shell color no longer changed, likely corresponding to a maximal bleaching of the shell. Several studies conducted on oysters and gastropods have also reported shell bleaching at pH ∼7.7-7.8, which could be attributed to delamination of the periostracum (Avignon et al., 2020; Duquette et al., 2017; Le Moullac et al., 2016; Sezer et al., 2018). Such alteration makes the shell more prone to CaCO_3_ dissolution (Peck et al., 2016; Tunnicliffe et al., 2009), leading to subsequent changes in biomineral microstructure and shell hardness (Auzoux-Bordenave et al., 2019; Chandra Rajan et al., 2021; Fitzer et al. 2014). Overall, we demonstrate rapid shell alterations due to pH reduction in both species. Compromised shell integrity, resulting from the periostracum delamination, is likely to impair animal fitness and make the oysters more vulnerable to predation and environmental stressors (Waldbusser et al., 2011; Wright et al., 2018).

### OA causes a major remodeling of membrane fatty acids

We show for the first time interspecific divergence in the remodeling of membrane FA in response to OA at pH_T_ 7.0 in *C. gigas* and pH_T_ 7.5 in *O. edulis*. Remodeling of membrane FA is pivotal to maintaining organism’s homeostasis in a changing environment (Lee et al., 2018). Similarly, Lutier et al. (2022) reported a major remodeling of FA at pH 7.0 in *C. gigas* juveniles. Overall, very few studies conducted on bivalves have investigated the effects of acidifying conditions on fatty acid composition and there is no such comparison in the literature in *O. edulis*.

Specifically, the level of DHA (22:6n-3) decreased with decreasing pH at the benefit of two fatty acids related to plasmalogenic phospholipids like NMI for *C. gigas* and DMA – a derivative product from plasmalogen transesterification – for *O. edulis*. These results are similar to those obtained in *C. gigas* by Lutier et al. (2022). We hypothesise that oysters’ membrane fatty acids were modified in response to low pH to protect cell membranes from oxidative stress while maintaining membrane fluidity. Indeed, DHA is a long-chain polyunsaturated fatty acid (PUFA) particularly susceptible to peroxidation (Munro and Blier, 2012), whereas NMI and plasmalogens can act as scavengers of reactive oxygen species (Barnathan, 2009; Kraffe et al, 2004; Leray et al, 2002; Munro and Blier, 2012). In addition, NMI and plasmalogens may contribute to maintaining membrane fluidity and ion exchanges between intra and extracellular compartments (Munro and Blier, 2012). Here, we show that the two oyster species use the same biochemical machinery to cope with low pH, but their tipping point vary markedly, suggesting different strategies.

### Oysters are tolerant to corrosive water

Both oyster species survive and grow at pH levels as low as 6.6 while seawater was undersaturated in calcite and aragonite (Ω_C_ = 0.2 and Ω_A_ = 0.1). Several studies have reported the ability of molluscs to calcify and grow in corrosive seawater, see e.g., the mussels *Mytilus edulis* in Berge et al. (2006) and *Bathymodiolus brevior* in Tunnicliffe et al. (2009), and the limpet *Patella caerulea* in Rodolfo-Metalpa et al. (2011).

Note however that the oysters were fed *ad libitum* and their energy reserves were unaffected by pH decrease. It is likely that food availability allowed oysters to cope with acidification while maintaining positive growth without depleting reserves. The effects of acidification on growth and energy may be more pronounced under limiting food conditions (Hettinger et al., 2013; Sanders et al., 2013; Thomsen et al., 2013). We may have underestimated the impacts of OA in our study and it would be interesting to obtain reaction norms under different food levels.

However, at pH_T_ > 6.6, oyster shells were smaller than at the onset of the experiment reflecting a net dissolution of the shell in both species. This coincides with a marked increase in mortality in *O. edulis* suggesting death by shell dissolution (Green et al., 2009). At this pH_T_, *C. gigas* showed no mortality and reserves were saved. We hypothesise that *C. gigas* went through metabolic depression to cope with such extreme acidification event. Indeed, organisms can shift to metabolic rate depression enabling it to delay the onset of homeostatic disturbances incompatible with survival (Guppy and Withers, 1999; Sokolova, 2021). This time-limited survival strategy was previously suggested in Lutier at al. (2022) in juvenile *C. gigas* exposed to extremely acidic waters thanks to additional respiration and calcification data. The absence of data on the state of reserves at this pH in *O. edulis* is inconclusive. Although both oyster species are tolerant to OA, they deploy different short-term survival strategies in highly corrosive waters, with *C. gigas* showing the greatest acclimation potential.

### Oyster species exhibit different physiological trade-offs to cope with OA

Reaction norms differed markedly between the two oyster species. Acidification has a greater effect on the growth of *C. gigas*, as shown by the steeper regression slopes, compared to *O. edulis*, but its survival is better. Under acidified conditions, *C. gigas* may allocate a greater proportion of energy to maintenance processes to stay alive at the expense of growth than *O. edulis*. Such trade-offs between growth and survival under stressful conditions was previously suggested in several studies (Gazeau et al., 2013; Stumpp et al., 2011; Thomsen and Melzner, 2010). In addition, the marked decrease in growth in *C. gigas* in response to low pH may reflect shell dissolution that could compensate for internal acidosis by increasing bicarbonate ions in tissues and extracellular fluids (Michaelidis et al., 2005). Conversely, *O. edulis* may allocate energy towards growth at the expense of homeostasis. A full energy budget is necessary to further investigate the physiological trade-offs in response to pH.

These different strategies exhibited by the two oyster species towards OA can be explained by their respective ecological niches. Intertidal organisms that live in highly variable environments are likely to be more tolerant to pH fluctuation than those living in the more stable subtidal zone (Melzner et al., 2009; Vargas et al., 2017, 2022). Comparative studies have suggested that the intertidal *C. gigas* could even outcompete the subtidal *O. edulis* under fluctuating environmental conditions (Gilson et al., 2021; Green et al., 2017; Stechele et al., 2022). Regarding to seawater acidification, Bamber (1990) established the reaction norms to pH in *C. gigas* and *O. edulis* after a 30-day exposure and reported the highest sensitivity of *O. edulis*, although the pH gradient and the experimental procedure for acidifying the water (adding sulphuric acid) may be questionable. Overall, our results show that the intertidal Pacific oyster may have a higher acclimation potential than the subtidal flat oyster to cope with rapid and extreme OA events, which corroborates the habitat-specific response hypothesis.

## Conclusion

Our study provides important data about the plasticity capacity and tolerance thresholds of two oyster species living in different habitats to future global changes. However, further works integrating several oyster populations at different life stages under different food regimes and temperatures are needed to fully characterise specific reaction norms. We emphasise the need for considering habitat-specific variability of pH/pCO_2_ based on *in situ* observations to design more realistic experiments.

## Acknowledgements

We thank the Ifremer staff from Argenton and Bouin for animal production and maintenance. We are particularly grateful to Moussa Diagne for phytoplankton production, Matthias Huber, Isabelle Queau and Jacqueline Le Grand for zootechnical support, Claudie Quere and Valerian Le Roy for biochemical analyses (Ifremer, LEMAR). We thank Vincent Ouisse and Elodie Foucault (Ifremer - Laboratoire Environnement Ressources Languedoc-Roussillon) for total alkalinity analyses.

## Author contributions

C.C., C.D.P. and F.P. designed and conducted the experiment. C.C. executed analyses with Ifremer staff. C.C. and M.L. performed statistical analyses. C.C. and C.D.P. wrote the original draft, and all authors contributed and approved the final manuscript. F.P. obtained the funding. This study is part of the PhD thesis of C.C.

## Funding

The study was financially supported by the Ocean Acidification Program of the French Foundation for Research on Biodiversity (FRB, www.fondationbiodiversite.fr) and the French Ministère de la transition écologique.

### Data availability

Raw data supporting the findings of this study are publicly available at the SEANOE Digital Repository: https://doi.org/10.17882/95793.

