## Supplemental Figures and Tables for "Differential reaction norms to ocean acidification in two oyster species from contrasting habitats"

**Supplemental file**

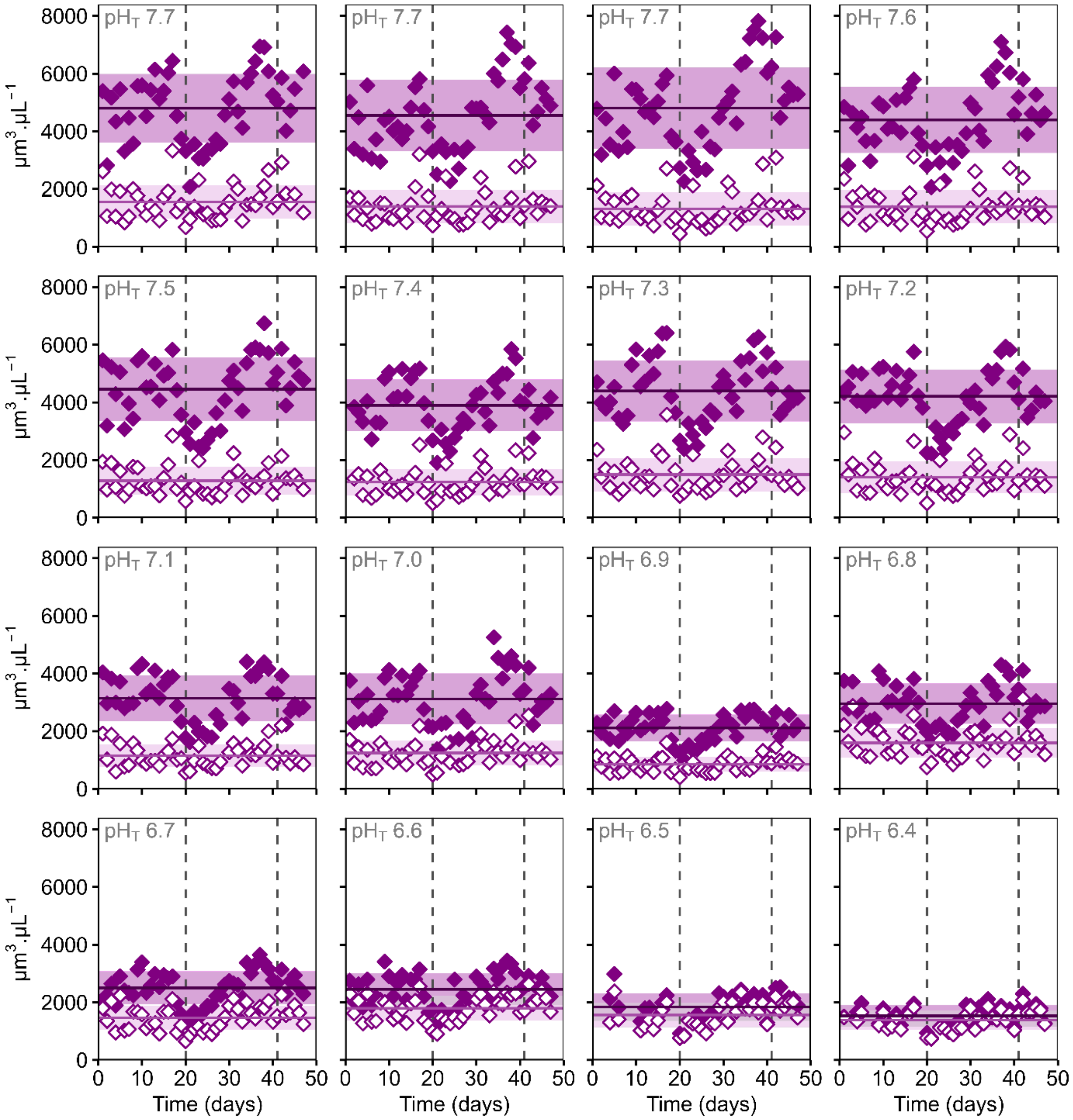


**Fig. S1. Temporal evolution of the phytoplankton cell volume at the tank inlet (filled symbols) and outlet (empty symbols) during 48 days of exposure under 16 $pH_T$ conditions.** This food consumption was quantified at the scale of the whole oyster batch in each tank as both *C. gigas* and *O. edulis* species were exposed in common garden. The horizontal lines correspond to the mean phytoplankton cell volume (± s.d.) over the entire experimental period. The vertical lines correspond to the sampling times of oysters, thus this decreased biomass causes a decrease in the food consumption right after. The mean phytoplankton concentration at the outlet of tanks was 1383 ± 214 $\mu m^3.\mu l^{-1}$. These are raw zootechnical data not standardized by weight.

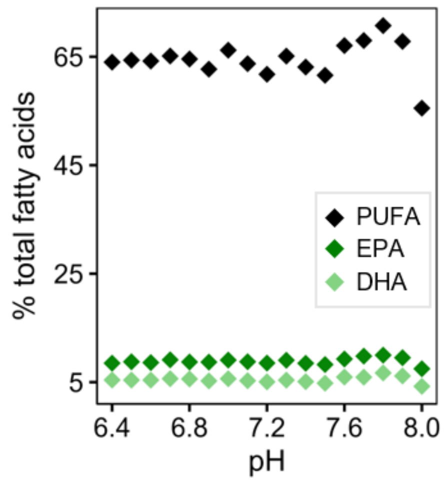


**Fig. S2. Essential fatty acid proportions in phytoplankton as a function of pH (on total scale).** Phytoplankton was exposed for two hours to 17 pH conditions considering a flow rate of 500 $ml.min^{-1}$. This experiment was performed after the main oyster experiment. Essential fatty acids were not correlated with pH decrease. Abbreviations: PUFA, total of long-chain polyunsaturated fatty acids (including EPA and DHA); EPA, eicosapentaenoic acid (20:5n-3); DHA, docosahexaenoic acid (22:6n-3).

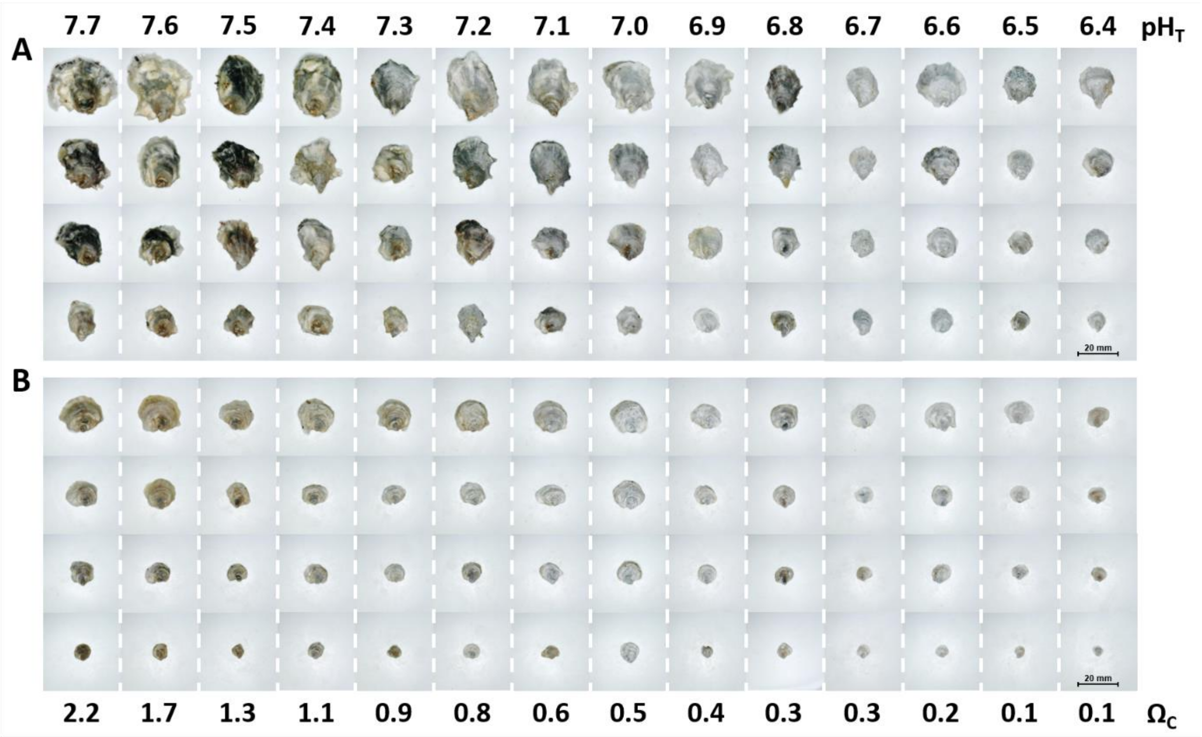


**Fig. S3. The gradual shell reduction and bleaching of *Crassostrea gigas* (A) and *Ostrea edulis* (B) after 20 days of exposure to $pH_T$ levels ranging from 7.7 to 6.4.** Only four oysters are shown from each condition and species, and a size variability is noted between individuals from the same condition. Corresponding seawater pH on the total scale ($pH_T$) and saturation state of calcite ($\Omega_C$) are indicated.

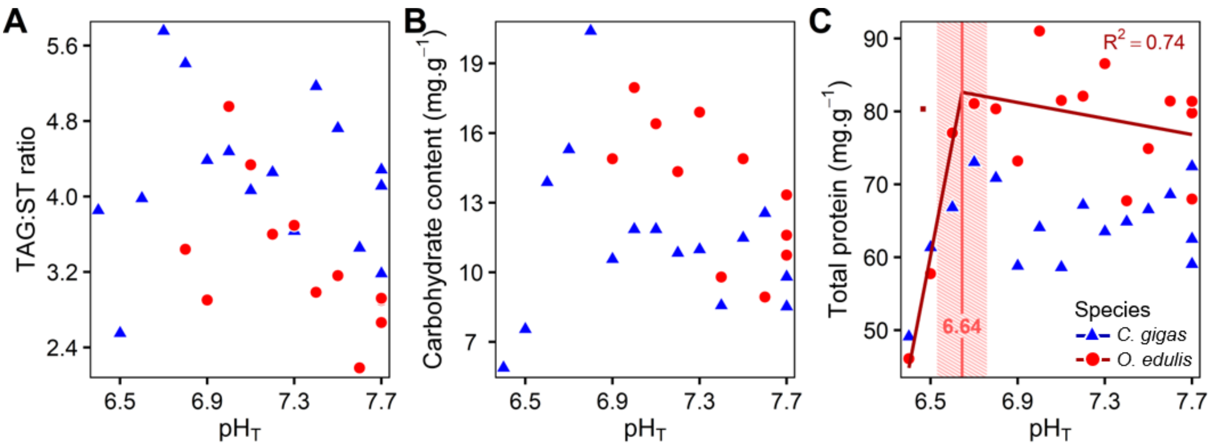


**Fig. S4. Energy reserves in *C. gigas* and *O. edulis* oysters as a function of pH (on total scale).** Triacylglycerol:sterol ratio (A), total carbohydrate content (B) and total protein content (C) in oysters after 41 days of exposure under 16 $pH_T$ conditions (n = 10 oysters pooled per condition and species). Tipping point and the 95% confidence interval of *O. edulis* is shown in striped red. The slopes were not significant ($p > 0.05$).

**Table S1. Relative contribution of membrane fatty acids in *Crassostrea gigas*.**

| Mean pHT | Fatty acid (% polar fatty acids) | | | | | | | | | |
|---|---|---|---|---|---|---|---|---|---|---|
| | 14:0 | 16:0 | 16:1n-7 | 18:0DMA | 18:0 | 18:1n-9 | 18:1n-7 | 18:2n-6 | 18:3n-3 | 18:4n-3 |
| 7.7 | 2.3 | 12.2 | 2.1 | 9.1 | 3.1 | 1.7 | 5.6 | 1.3 | 1.1 | 1.6 |
| 7.7 | 2.2 | 12.2 | 1.9 | 9.3 | 3.9 | 1.7 | 5.2 | 1.3 | 1.2 | 1.6 |
| 7.7 | 2.2 | 11.9 | 1.9 | 9.1 | 3.6 | 1.8 | 5.3 | 1.4 | 1.3 | 1.6 |
| 7.6 | 2.3 | 11.9 | 2.0 | 9.1 | 3.1 | 1.7 | 5.9 | 1.3 | 1.1 | 1.6 |
| 7.4 | 2.4 | 11.2 | 2.1 | 8.9 | 3.1 | 1.8 | 5.8 | 1.4 | 1.2 | 1.7 |
| 7.3 | 2.3 | 12.0 | 2.0 | 7.0 | 3.1 | 1.9 | 6.0 | 1.5 | 1.3 | 2.0 |
| 7.2 | 2.1 | 11.1 | 1.9 | 8.9 | 3.4 | 1.8 | 5.9 | 1.4 | 1.2 | 1.9 |
| 7.1 | 1.9 | 10.1 | 1.8 | 9.7 | 3.7 | 1.8 | 5.6 | 1.5 | 1.3 | 1.9 |
| 7.0 | 1.9 | 10.5 | 1.6 | 9.5 | 3.9 | 1.8 | 5.5 | 1.3 | 1.3 | 1.7 |
| 6.9 | 2.0 | 11.1 | 1.9 | 9.0 | 3.5 | 1.8 | 5.9 | 1.4 | 1.1 | 1.7 |
| 6.8 | 2.0 | 11.4 | 2.1 | 7.9 | 4.5 | 1.9 | 6.3 | 1.3 | 1.2 | 1.6 |
| 6.7 | 2.0 | 10.5 | 2.2 | 8.3 | 3.5 | 1.9 | 6.8 | 1.4 | 1.1 | 1.7 |
| 6.6 | 1.4 | 8.8 | 2.0 | 8.7 | 4.0 | 1.7 | 6.2 | 1.0 | 0.9 | 1.1 |
| 6.5 | 1.4 | 9.4 | 2.1 | 7.7 | 4.1 | 2.0 | 6.2 | 0.9 | 0.7 | 1.0 |
| 6.4 | 1.0 | 9.6 | 1.8 | 9.0 | 4.9 | 2.4 | 5.6 | 0.6 | 0.3 | 0.5 |
| Contribution to PC1 (%) | 4.5 | 28.9 | 0.0 | 1.5 | 4.7 | 0.4 | 1.1 | 1.3 | 1.4 | 3.0 |
| Correlation to PC1 | 0.9 | 0.9 | -0.2 | 0.3 | -0.7 | -0.7 | -0.4 | 0.8 | 0.8 | 0.8 |

| Mean pHT | Fatty acid (% polar fatty acids) | | | | | | | | |
|---|---|---|---|---|---|---|---|---|---|
| | 20:1DMA | 20:0 | 20:1n-11 | 20:1n-7 | 20:4n-6 | 20:5n-3 | $22{:}2_{i,j}$NMI | 22:5n-6 | 22:6n-3 |
| 7.7 | 2.3 | 1.3 | 1.4 | 4.0 | 3.9 | 11.8 | 5.1 | 3.0 | 17.1 |
| 7.7 | 1.8 | 1.2 | 1.8 | 4.0 | 4.0 | 10.6 | 5.2 | 3.3 | 17.8 |
| 7.7 | 1.9 | 1.3 | 1.9 | 4.2 | 3.8 | 10.7 | 5.5 | 3.1 | 17.7 |
| 7.6 | 2.0 | 1.3 | 1.5 | 4.0 | 3.9 | 12.2 | 4.9 | 3.1 | 17.6 |
| 7.4 | 2.1 | 1.1 | 1.6 | 4.1 | 3.8 | 12.2 | 5.1 | 3.1 | 17.0 |
| 7.3 | 1.6 | 1.1 | 1.6 | 4.3 | 3.9 | 12.8 | 5.2 | 3.1 | 17.6 |
| 7.2 | 1.9 | 1.1 | 1.6 | 4.1 | 4.0 | 12.0 | 5.3 | 3.1 | 17.9 |
| 7.1 | 1.9 | 1.1 | 1.8 | 4.2 | 4.0 | 11.3 | 5.5 | 3.2 | 17.9 |
| 7.0 | 2.0 | 1.2 | 1.9 | 4.5 | 4.0 | 11.2 | 6.0 | 3.0 | 17.7 |
| 6.9 | 2.1 | 1.5 | 1.6 | 4.3 | 4.0 | 11.8 | 5.4 | 2.9 | 17.4 |
| 6.8 | 1.8 | 1.1 | 1.7 | 4.6 | 4.2 | 12.3 | 5.5 | 2.4 | 15.5 |
| 6.7 | 2.0 | 1.2 | 1.6 | 4.5 | 4.0 | 12.3 | 5.9 | 2.5 | 16.2 |
| 6.6 | 2.7 | 1.1 | 2.1 | 5.2 | 4.9 | 12.4 | 7.3 | 2.2 | 16.0 |
| 6.5 | 2.7 | 1.1 | 1.7 | 5.3 | 4.9 | 12.7 | 7.0 | 2.1 | 16.1 |
| 6.4 | 3.3 | 1.7 | 1.6 | 5.3 | 3.9 | 12.8 | 6.5 | 2.1 | 15.7 |
| Contribution to PC1 (%) | 3.9 | 0.0 | 0.1 | 6.7 | 2.0 | 5.7 | 13.3 | 5.0 | 16.5 |
| Correlation to PC1 | -0.8 | -0.1 | -0.3 | -1.0 | -0.7 | -0.6 | -0.9 | 1.0 | 0.8 |

The relative contribution (%) of each fatty acid is indicated for the 16 $pH_T$ conditions. Only fatty acid contributing to >1% were considered. Their contribution and correlation to the first principal component (PC1) of the PCA (Fig. 5A) are indicated at the bottom of the table.

### Table S2. Relative contribution of membrane fatty acids in *Ostrea edulis*.

| Mean pHT | Fatty acid (% polar fatty acids) | | | | | | | | | |
|---|---|---|---|---|---|---|---|---|---|---|
| | 14:0 | 16:0 | 16:1n-7 | 18:0DMA | 18:0 | 18:1n-9 | 18:1n-7 | 18:2n-6 | 18:3n-3 | 18:4n-3 |
| 7.7 | 1.8 | 9.1 | 2.1 | 11.3 | 4.1 | 1.9 | 2.9 | 1.5 | 1.2 | 1.0 |
| 7.7 | 2.0 | 9.3 | 2.1 | 11.4 | 4.6 | 2.0 | 3.0 | 1.6 | 1.4 | 1.2 |
| 7.7 | 2.1 | 9.9 | 2.2 | 10.8 | 4.0 | 2.1 | 3.0 | 1.6 | 1.5 | 1.1 |
| 7.6 | 1.9 | 9.4 | 2.2 | 10.5 | 4.1 | 2.0 | 3.1 | 1.5 | 1.1 | 1.1 |
| 7.5 | 2.0 | 9.7 | 2.3 | 10.3 | 4.2 | 2.1 | 3.3 | 1.7 | 1.3 | 1.2 |
| 7.4 | 1.7 | 8.6 | 2.1 | 10.3 | 4.0 | 1.9 | 3.9 | 1.5 | 1.2 | 1.3 |
| 7.3 | 1.9 | 9.6 | 2.2 | 11.5 | 4.3 | 2.0 | 3.1 | 1.5 | 1.1 | 1.1 |
| 7.2 | 1.7 | 9.5 | 2.1 | 11.8 | 4.3 | 1.7 | 2.9 | 1.4 | 1.0 | 1.0 |
| 7.1 | 1.6 | 9.1 | 2.0 | 11.2 | 4.6 | 1.8 | 3.0 | 1.4 | 1.3 | 1.0 |
| 7.0 | 1.6 | 9.5 | 2.1 | 12.0 | 4.7 | 1.8 | 2.9 | 1.3 | 1.2 | 1.0 |
| 6.9 | 1.5 | 9.2 | 2.0 | 11.6 | 4.3 | 1.7 | 2.9 | 1.3 | 1.1 | 1.0 |
| 6.8 | 1.4 | 9.4 | 1.8 | 11.3 | 5.1 | 1.9 | 2.7 | 1.4 | 1.1 | 1.1 |
| Contribution to PC1 (%) | 3.7 | 0.0 | 0.9 | 29.7 | 6.8 | 1.3 | 6.6 | 1.3 | 0.8 | 1.4 |
| Correlation to PC1 | 0.7 | -0.0 | 0.7 | -0.8 | -0.7 | 0.7 | 0.7 | 0.8 | 0.5 | 0.8 |

| Mean pHT | Fatty acid (% polar fatty acids) | | | | | | | | |
|---|---|---|---|---|---|---|---|---|---|
| | 20:1DMA | 20:0 | 20:1n-11 | 20:1n-7 | 20:4n-6 | 20:5n-3 | $22:2_{i,j}$NMI | 22:5n-6 | 22:6n-3 |
| 7.7 | 1.2 | 0.7 | 1.5 | 5.1 | 5.1 | 13.9 | 7.2 | 3.2 | 15.2 |
| 7.7 | 1.1 | 0.6 | 1.4 | 4.8 | 4.6 | 13.1 | 6.8 | 3.4 | 16.0 |
| 7.7 | 1.1 | 0.8 | 1.3 | 4.8 | 4.8 | 13.2 | 6.8 | 3.3 | 15.2 |
| 7.6 | 1.2 | 0.8 | 1.4 | 5.0 | 5.0 | 14.2 | 7.1 | 3.1 | 14.9 |
| 7.5 | 1.0 | 0.7 | 1.3 | 4.7 | 4.9 | 13.3 | 6.5 | 3.2 | 16.1 |
| 7.4 | 1.4 | 0.6 | 1.3 | 5.0 | 5.0 | 14.4 | 6.6 | 2.9 | 16.1 |
| 7.3 | 1.2 | 0.7 | 1.2 | 5.0 | 4.8 | 13.3 | 6.9 | 3.1 | 15.5 |
| 7.2 | 1.2 | 0.8 | 1.3 | 5.2 | 4.9 | 14.2 | 6.8 | 2.9 | 15.3 |
| 7.1 | 1.1 | 0.7 | 1.4 | 5.1 | 5.0 | 14.0 | 7.1 | 2.8 | 15.2 |
| 7.0 | 1.1 | 0.8 | 1.4 | 4.8 | 4.8 | 14.1 | 6.7 | 2.6 | 14.8 |
| 6.9 | 1.2 | 0.8 | 1.3 | 5.3 | 5.3 | 14.4 | 7.1 | 2.6 | 14.9 |
| 6.8 | 1.0 | 0.7 | 1.4 | 5.0 | 4.9 | 13.2 | 7.2 | 2.4 | 14.2 |
| Contribution to PC1 (%) | 0.2 | n.a. | 0.2 | 1.2 | 0.5 | 1.6 | 3.1 | 7.8 | 33.0 |
| Correlation to PC1 | 0.4 | n.a. | -0.4 | -0.5 | -0.3 | -0.2 | -0.6 | 0.8 | 0.9 |

The relative contribution (%) of each fatty acid is indicated for the 16 $pH_T$ conditions. Only fatty acid contributing to >1% were considered (then not 20:0). Their contribution and correlation to the first principal component (PC1) of the PCA (Fig. 5B) are indicated at the bottom of the table.
